# Proximate Predictors of Variation in Egg Rejection Behavior by Hosts of Avian Brood Parasites

**DOI:** 10.1101/788505

**Authors:** Mikus Abolins-Abols, Mark E. Hauber

**Author notes:** Correspondence should be addressed to Mikus Abolins-Abols. Funding for this project was provided by the Harley Jones Van Cleave Professorship to MEH. We thank private landowners for their permission to work on their properties.

## Abstract

The rejection of parasitic eggs by hosts of avian brood parasites is one of the most common and effective defenses against parasitism. Despite its adaptive significance, egg rejection often shows substantial intraspecific variation: some individuals are more likely to remove or abandon parasitic eggs than others. Understanding variation in egg rejection requires that we study factors linked to both the ability to perceive parasitic eggs, as well as factors that may influence the rejection of a foreign egg once it has been recognized. Here we asked what cognitive, physiological, and life-history factors explained variation in the rejection of model eggs by American Robin *Turdus migratorius* females. We found that the probability of egg rejection was related to the clutch size at the time of parasitism: in support of Weber’s law, females with fewer eggs were more likely to reject the model eggs. In turn, females with greater mass and higher corticosterone levels were less likely to reject eggs, and egg rejection probability was negatively related to incubation progress. Our data thus suggest that proximate predictors of an individual’s egg rejection behavior include components of the nest’s perceptual environment, life-history factors, as well as the physiological state of the animal. However, much of the variation in the responses of robins to the model eggs remained unexplained. Future experiments should aim to understand the causal roles of these and other factors in generating within- and among-individual variation in the rejection of parasitic eggs.

## Introduction

In obligate avian brood parasitic systems, parasites lay their eggs in the nests of host species, often causing a substantial reproductive loss to the affected hosts (Davies, 2000; Feeney, Welbergen, & Langmore, 2014; Soler, 2017). Many hosts have evolved a variety of adaptations that allow them to mitigate or overcome the negative fitness effects of brood parasitism. One of the most effective defenses against parasitism is the rejection of foreign eggs from the nest (Davies & Brooke, 1988; Moksnes & RØskaft, 1992; Rothstein, 1982). Decades of research have demonstrated that some hosts can recognize parasitic eggs using a variety of cognitive rules, which include memory-based templates as well as direct comparisons of egg colors (reviewed by Manna, Moskát, & Hauber, 2017). Brood parasites, on the other hand, have evolved to overcome these defenses by laying mimetic (Davies & Brooke, 1988) or cryptic (Langmore, Stevens, Maurer, & Kilner, 2009) eggs that limit the ability of hosts to recognize and reject the parasitic eggs (Davies, Brooke, & Kacelnik, 1996; Feeney, Welbergen, & Langmore, 2012; Moskát & Hauber, 2007).

Variation in the host visual systems, morphology, and coevolutionary history with brood parasites has led to divergent abilities of host species to recognize and reject parasitic eggs (Avilés, 2008; Peer & Sealy, 2004; Rothstein, 1975; Stoddard & Stevens, 2011). Importantly, however, differences in the egg rejection behavior exist not only among, but also within species (Croston & Hauber, 2014a; Grim, Samaš, & Hauber, 2014; Molina-Morales et al., 2014; Samaš, Hauber, Cassey, & Grim, 2011). While some of the intraspecific variation in egg rejection is linked to the variation in the degree of mimicry of the parasitic egg in a particular clutch (Abolins-Abols, Hanley, Moskát, Grim, & Hauber, 2019; Moskát et al., 2014), host individuals nevertheless often show variation in the response to the *same* foreign egg stimuli (Grim et al., 2014; Luro & Hauber, 2017). From a mechanistic standpoint, differences in responses to the same parasitic stimuli could be caused by differences in the ability of hosts to perceive the parasitic egg stimulus, or differences in the probability of rejection once the parasitic egg has been recognized.

Differences in the ability to perceive and identify a parasitic egg as non-self may be driven either by individual variation in sensory systems (Stoddard & Stevens, 2011) or variation of the perceptual environment of the parasitized nest (Honza, Procházka, Morongová, Capek, & Jelínek, 2011). For example, some hosts use the range and distribution of egg colors and patterns in the nest to make rejection decisions (Bán, Moskát, Barta, & Hauber, 2013; Moskát, Avilés, Bán, Hargitai, & Zölei, 2008). The variation of egg color in a clutch may be driven by the extent of the host’s intraclutch variability (Moskát et al., 2008) or differences in the parasite-to-host egg ratio, due to single vs. multiple parasitism and/or variation in host clutch size (Manna et al., 2019; Moskát et al., 2009; Stevens, Troscianko, & Spottiswoode, 2013, but see Lang, Bollinger, & Peer, 2014). Hosts may respond differently to nests with varying parasite-to-host egg ratio not only because of the variation in color, but also because of the Weber’s law, which posits that the ability of animals to discriminate between stimuli depends on the stimulus proportionality (Stevens, 1975). Weber’s law states that, when the proportional change in a stimulus is low (e.g. a single parasitic egg added to a large clutch), the ability of individuals to detect that change is lower relative to situations when the proportional change is larger (e.g. a single parasitic egg in a small clutch). The ability of hosts to recognize brood parasitism may thus depend on the size of the host clutch (Akre & Johnsen, 2014).

The probability of parasitic egg rejection may depend not only on the variation in the host’s ability to perceive the parasitic egg, but also on factors that affect the likelihood of egg rejection *after* sensory and cognitive processes have processed the parasitic egg stimulus. For example, hosts are more likely to reject foreign eggs of the same phenotype following a prior experience with brood parasitism (Hauber, Moskát, & Bán, 2006) or earlier in the incubation stage (Moskát & Hauber, 2007). Such experience- and life history-dependent plasticity in parasitic egg rejection reflects the complexity of the changing costs associated with brood parasitic egg rejection in relation to host fitness (Avilés, 2018; Hauber et al., 2014; Molina-Morales et al., 2014). For example, a brood parasitic chick hatched from an egg laid in the host nest prior to the onset of incubation is more likely to negatively affect host fitness than a parasitic chick hatched from an egg laid late in incubation (Moskát, 2005). Variation in the likelihood of egg rejection across different life history stages may in turn be linked to changes in the physiological state of the host (Abolins-Abols & Hauber, 2018): for instance, hormones that affect maternal motivation and attachment in females (Bridges, 2015; Richard-Yris, Leboucher, Chadwick, & Garnier, 1987) may cause changes in the readiness of hosts to accept or reject any eggs in the nest (Abolins-Abols & Hauber, 2018). Specifically, corticosterone, a glucocorticoid hormone that is involved in the regulation of the stress-response and metabolism (MacDougall-Shackleton, Bonier, Romero, & Moore, 2019; Romero & Butler, 2007) can suppress maternal behavior (Horton & Holberton, 2009). Importantly, corticosterone has been shown to increase in hosts when they are exposed to brood parasitism (Mark & Rubenstein, 2013; Ruiz-Raya et al., 2018), suggesting that it may affect the likelihood of parasitic egg rejection by hosts (Abolins-Abols & Hauber, 2018). In addition to context-specific, plastic differences in egg rejection, hormones may also explain stable and repeatable individual variation (i.e., personalities) in egg rejection irrespective of the context (Avilés & Parejo, 2011). Indeed, in many cases hosts show repeatable individual variation in parasitic egg rejection (Croston & Hauber, 2014a; Samaš et al., 2011) often across different contexts (Grim et al., 2014). Importantly, hormonal levels also often show consistent differences between individuals (Baugh et al., 2017; Rensel & Schoech, 2011; Romero & Reed, 2008), suggesting that individual differences in egg rejection may be hormonally mediated (Abolins-Abols & Hauber, 2018).

In this study, we took a holistic approach and asked what cognitive, physiological, and life-history factors explained variation in the egg rejection behavior of American Robins (hereafter: robin), hosts to the obligate brood-parasitic Brown-headed Cowbird *Molothrus ater* (hereafter: cowbird) in North America. Robins have served as a productive model system in which to investigate the perceptual mechanisms underlying parasitic egg rejection (e.g. Hanley et al., 2017). While only a minority of robin individuals accept naturally laid parasitic cowbird eggs (Rothstein, 1975, 1982), they show repeatable individual variation in the acceptance or rejection of model eggs (Croston & Hauber, 2014a; Luro & Hauber, 2017).

Psychophysical theory (Weber’s law; Akre & Johnsen, 2014) and empirical data (Moskát et al., 2008) suggest that hosts should reject parasitic eggs more frequently when they have smaller clutches. We therefore asked whether the probability of rejection of model parasitic eggs by robins was explained by clutch size at the time of experimentation. We also tested if egg rejection was related to the life history context (Hauber et al., 2014) by assessing whether model egg rejection by robin hosts depended on female age, the progression of the breeding season, and incubation stage. We predicted that older, more experienced breeders may be more likely to reject model eggs (Moskát, Bán, & Hauber, 2014), that females would be more likely to accept model eggs added later in incubation (Alvarez, 1996; Moskát, 2005) and later in the breeding season (Thorogood & Davies, 2013). Finally, we asked if the probability of egg rejection depended on host internal state as assessed by baseline corticosterone levels and body mass, predicting that females with higher corticosterone levels (Abolins-Abols & Hauber, 2018) and higher body mass (Peer & Sealy, 2004) would be more likely to reject model eggs.

## Method

### Study Site and Species

All procedures were approved by the University of Illinois IACUC protocol #17259. We studied American Robin females at Wandell’s Nursery, a deciduous tree farm near Urbana, IL, USA (lat: 40.128184; long: −88.105349), between April 26 and July 8, 2018. We surveyed the study area every 3 days to search for nests and monitor their status. We assumed a deposition of one new egg per day during the laying period, and clutches with 2 or more consecutive days of constant egg numbers were deemed completed (Vanderhoff, Pyle, Patten, Sallabanks, & James, 2016). The median clutch size is 3 eggs in this population (mean: 3.38, range: 2-5).

### Capture and Sample Collection

We captured female robins (n=52) using mist nets as close to the predicted clutch completion date as possible (range: 2 days before, 4 days after; median 1.5 days after completion). In bird species with female-only incubation, including the American Robin (Vanderhoff et al., 2016), the female is known to be the egg rejecter sex (Honza, Požgayová, Procházka, & Tkadlec, 2007; Palomino, Martín-vivaldi, Soler, & Soler, 1998). Subjects were captured between 6:00 am and 11:30 am using a 12 m long mist net (eye size 38 mm) that was bent around a middle pole, resulting in a V-shaped net around the focal female’s nest. Robin females were either caught in the net while attempting to land in their nests, or, if the females avoided the net and started incubating, by flushing the incubating females into the net.

Upon capture, we collected a ∼75 µl blood sample from the brachial vein using a 26 G needle and heparinized capillary tubes. The mean start time of blood collection was 131 s after capture (range 70-220 s, standard deviation (SD) = 37.7). We typically ended blood collection at 180 s (3 min) after capture if a sufficient blood volume was collected (i.e., the “3 minute rule” for baseline corticosterone analysis: Romero & Reed, 2005). If not, we continued to collect blood after the 3 min cutoff (mean: 200 s, range for ending blood collection 143-245 s, SD = 18.9; see below for the effect of the timing of blood collection on corticosterone levels). Blood was transferred to a centrifuge tube and stored on ice until centrifugation (3-9 hrs after collection). We centrifuged samples at 13,000 rpm for 10 min at 4 °C to separate plasma from blood cells, and stored the plasma samples at −80 °C until analysis.

After blood collection, we banded each bird with a USGS aluminum band as well as 3 unique plastic color bands for individual identification in the field. We measured the mass of birds to the nearest 0.5 g. We estimated the age of the females (second year or after second year) by comparing the coloration of the wing feathers according to published guidelines (Pyle, 1997). Birds were then released at the site of capture (∼10 min after capture), and the net was removed.

One day after the capture, we returned to the focal nest to confirm female identity and nest attendance by verifying the band colors. We found that 6 out of 52 (12 %) of females abandoned their nests overnight and 5 out of 52 (10%) nests were depredated overnight following the capture.

### Experimental Parasitism

If the nest was still active on the day after capture, we placed a dark blue 3D-printed model cowbird egg in the nest (Figure 1). Our 3D-printed smooth nylon eggs (sourced commercially from Shapeways.com) resembled natural Brown-headed Cowbird eggs in size and weight (details given in Igic et al., 2015). We painted the eggs with acrylic dark blue paint (Winsor & Newton Galeria Acrylic Ultramarine) applying three paint coats to each egg. The resulting reflectance spectra of the model eggs and those of natural robin eggs have been reported previously (Croston & Hauber, 2014b). We did not remove a robin egg from the nest as cowbirds do when they parasitize nests (Davies, 2000), because prior studies had suggested that *Turdus* thrushes respond to model parasitic eggs similarly, irrespectively of whether clutch size is maintained or increased as part of the experiment (Grim et al., 2011). We chose this artificial egg color type because cowbirds are natural brood parasites of robins (Rothstein, 1982) and because eggs of this color have been shown to be rejected at intermediate rates by robin females (Croston & Hauber, 2014b). Furthermore, differences in the propensity of rejection or acceptance of these eggs are significantly repeatable among robin individuals across multiple exposures within a breeding attempt (Luro & Hauber, 2017).

**Figure 1.**
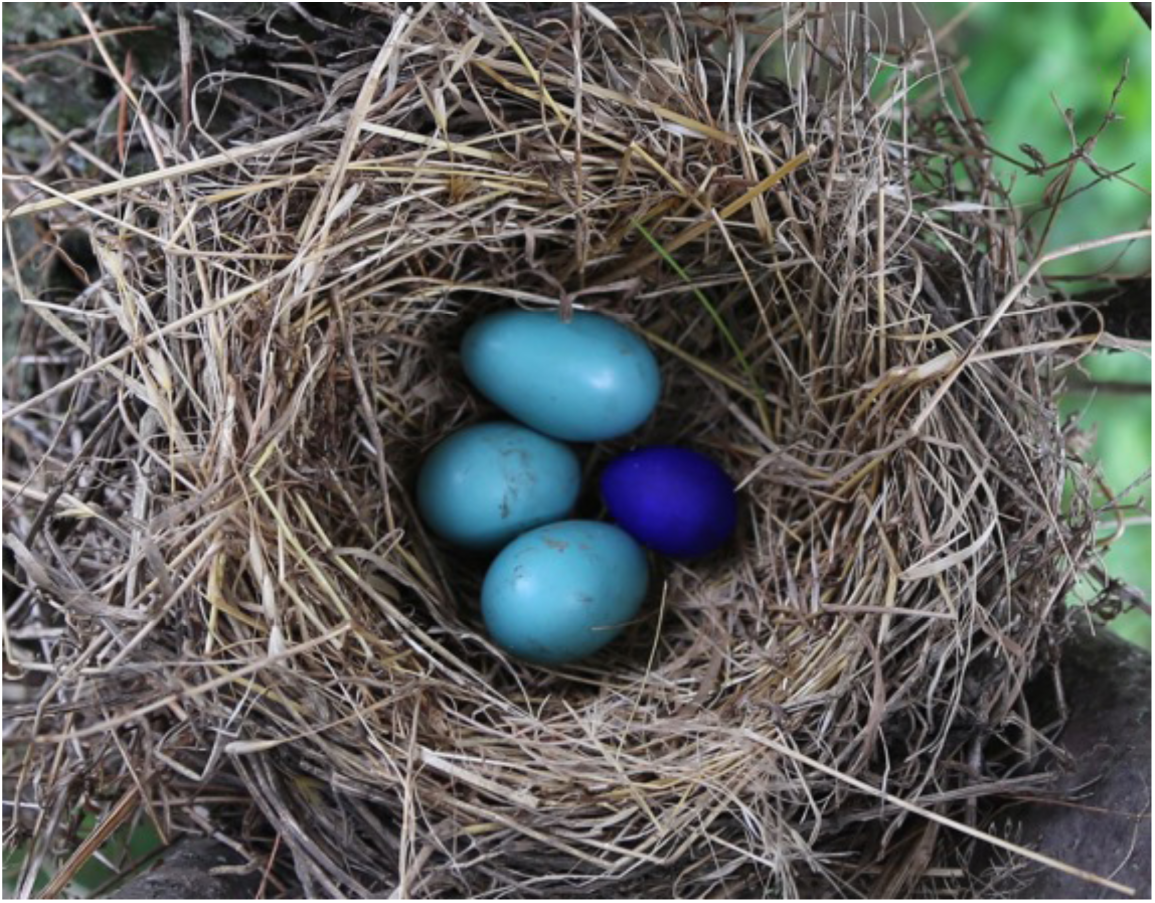
Model egg (dark blue, right) alongside three natural American Robin eggs (light blue). Photo by MAA.

Following the addition of the model egg, we counted the number of natural robin eggs in the nest. We then returned and recorded the presence/absence of the blue egg on the next day, visiting the nest at the same time of day compared to when the model egg was added. Robins that reject dark blue model eggs nearly always do so on the first day after experimental parasitism (Croston & Hauber, 2014a), therefore we used the rejection at day 1 as our response variable. We found that a further 3 nests were abandoned and 4 depredated overnight following the experimental parasitism. Due to the abandonment and predation of nests, and because 3 birds escaped during the initial processing, we had complete data on the rejection behavior of 31 birds (60% of the initial captures).

Because our aim was to begin to explore correlates of individual variation in rejection behavior in response to a model egg of known intermediate rejection rate, we did not manipulate any of the factors that were predicted to explain variation in egg rejection. We also did not investigate the difference in the rejection rates of the dark blue model eggs in comparison with control model egg treatments, such as robin-mimetic or cowbird-mimetic colors: previous work had already shown these other egg colors to be consistently rejected or accepted, respectively, independent of the individual identity in robins (Croston & Hauber, 2014b; Hanley et al., 2017; Igic et al., 2015). However, as an internal control, we checked on each of the robin’s own eggs in the nest (i.e. their number and whether intact) throughout our monitoring period, and found none of these eggs to be missing or broken, indicating that robins responded specifically to the dark blue model eggs.

### Hormone Assays

We used an enzyme immune assay to measure the circulating baseline corticosterone levels in blood samples from females caught before the experimental parasitism (Cayman Chemical, catalogue number 501320). Hormones were extracted from plasma using diethyl ether: 10 µl plasma was suspended in 200 µl double-distilled water and mixed with 1 ml diethyl ether. The aqueous phase was flash-frozen, and the ether phase decanted. The extraction procedure was repeated 3 times, following which the ether was evaporated under a gentle stream of nitrogen using an in-house constructed evaporation manifold (Nevins et al., 2005). This extraction technique has been shown by MA-A to have high extraction efficiency for songbird plasma (96%, Abolins-Abols, Hope, & Ketterson, 2016). Therefore, to limit the logistical challenges associated radioactivity-based efficiency measurements, we did not measure extraction efficiency in this study (Virgin & Rosvall, 2018).

The plasma extract was suspended in 600 µl assay buffer provided with the kit by vortexing for 1 min before storing it at 4 °C overnight. The corticosterone concentration of the extract was measured following the instructions from the kit manufacturer. We first validated the Cayman corticosterone assay to determine that robin plasma extracts reacted with the assay as expected. The assay showed good parallelism (slope=1.00; r^2^=0.99), sensitivity (10.7 pg/mL), and recovery (88.9%). We then ran the extracted samples in triplicate, using a pooled robin plasma extract as a within- and across-plate control. We used a Biotek 800TS plate reader to record assay absorbance at 405 nm. Data were analyzed using a 4-point logistic curve using the analysis spreadsheet provided by Cayman Chemical. All samples fell within 20-80% B/Bo range. The among-plate coefficient of variation (CV) was 16.4%, whereas the within-plate CV was 5.17%. We averaged the data from the 3 replicates, and standardized sample concentrations across plates using the mean concentration of the control samples.

### Data Analysis

We first investigated if corticosterone levels in robin females were related to the timing of capture and blood sampling. While in most animals corticosterone levels start rising after 3 min, and therefore a blood sample taken under 3 min post capture can be considered a “baseline” sample (Romero & Reed, 2005), in some cases corticosterone levels start rising earlier (Small et al., 2017). In our data, corticosterone levels were positively correlated with the blood collection start time - circulating corticosterone in robins started to increase soon after capture (estimate=0.02 ng/ml increase per s; r^2^=0.16; p=0.02, Figure S1). We therefore corrected for the variation in corticosterone as a function of the blood collection start time by running a linear regression between these variables, and used the residuals as a bleed time-independent measure of baseline corticosterone levels. We chose to use the blood collection start time as opposed to the end time, because in the majority of cases we ended blood collection at exactly 3 min. No other covariates (time of day, time since nest approach, time taken to set up the mist net) were related to corticosterone levels (all p > 0.05, data not shown).

We used generalized linear models (GLMs) with binomial error structure in R (R Core Team, 2017) to ask if rejection of the model eggs was related to the following ecological and proximate factors, all of which were known to be linked to egg rejection in brood parasite hosts, including robins: the date of model egg addition (Dainson, Hauber, López, Grim, & Hanley, 2017), age (Moskát et al., 2014), mass (Peer & Sealy, 2004), incubation stage relative to clutch completion (further as “days since clutch completion”; Marchetti, 2000), the number of eggs in the nest at the time of experimentation (Moskát & Hauber, 2007), and baseline corticosterone (Ruiz-Raya et al., 2018). We did not have any *a priori* expectations for interactions between these variables. All predictors were standardized to mean 0 and standard deviation of 1.

To investigative which of our observed predictor variables best explained egg rejection we used information-theoretic (IT) approach, combined with model averaging (Burnham, Anderson, & Huyvaert, 2011; Symonds & Moussalli, 2011). The emerging consensus for animal behavior-relevant studies is that IT-based model averaging approaches are better suited for hypothesis testing with correlative datasets from natural study systems compared to the more traditional p-value based approaches (Aho, Derryberry, & Peterson, 2014). In short, an IT-based approach allows one to compare the strength of evidence for various competing alternative models that may explain the variation in a variable (here: egg rejection), while a frequentist p-value-based approach is a better fit for controlled experiments with a defined null hypothesis (Murtaugh, 2014; Valpine, 2014). An IT approach is especially useful when, as is the case in our data, there are a number of independent candidate predictors that may explain variation the response variable, but where a model with all of the covariates may not necessarily generate the best model. Model averaging of competing models, in turn, allows one to evaluate the strength of evidence supporting a relationship between a particular independent variable and the dependent variable (here, the probability of egg rejection). Nevertheless, we have also provided p-values for the variables included in the best models to aid in data interpretation.

We used the package *MuMIn* (Bartoń, 2018) to calculate the corrected Akaike information criterion (AICc) scores for the models that contained all combinations of predictors (without interaction terms) to determine the model that was most likely to be the best model. Ten models had AICc scores higher than the null model. We calculated the goodness of fit of each of these top models in relation to the null model using the package *lmtest* (Zeileis & Hothorn, 2002). To estimate how much variance the top models explained, we used the package *pscl* (Jackman, 2017) to calculate McFadden’s pseudo-R^2^. Seven models had AICc scores that were within 2.0 units of the top model, indicating a level of uncertainty of which variables contributed to the best model. We, therefore, also followed a model averaging approach (Burnham et al., 2011; Symonds & Moussalli, 2011): we calculated the relative importance of each independent variable within the models with cumulative AICc weights of 0.95, i.e. a set that had a 95% probability of including the best model.

## Results

Among the 31 female American Robins that were included in the analyses, 18 (58.1%) ejected the model egg within 24 hrs.

Out of candidate 64 statistical models testing the predictors of the probability of egg rejection, 10 had AICc-s higher than the null model, which included no predictor variables (Table 1). However, these models were highly competitive, with seven top models being within 2.0 AICc-s of each other. The seven top models included egg number, days since clutch completion, corticosterone levels, and mass as the predictors of egg rejection. The model with the lowest AICc (41.80) suggested that females with more eggs in the nest at the time of experimentation were significantly less likely to reject the model egg (Figure 2). However, based on the corrected Akaike weights (w_i_=0.11), this model only had 11% overall probability of being the best model for explaining egg rejection behavior, and it explained only 11% of overall variation in egg rejection (r^2^=0.11). An equally competitive model (AICc=42.75, ΔAICc=0.95, w_i_=0.07, r^2^=0.28), included all of the covariates present in the 7 top models, and suggested that females with smaller clutch sizes, lower mass, lower corticosterone levels, and those whose nests were parasitized earlier with respect to their clutch completion were more likely to reject the model eggs (Table 2).

**Table 1.**
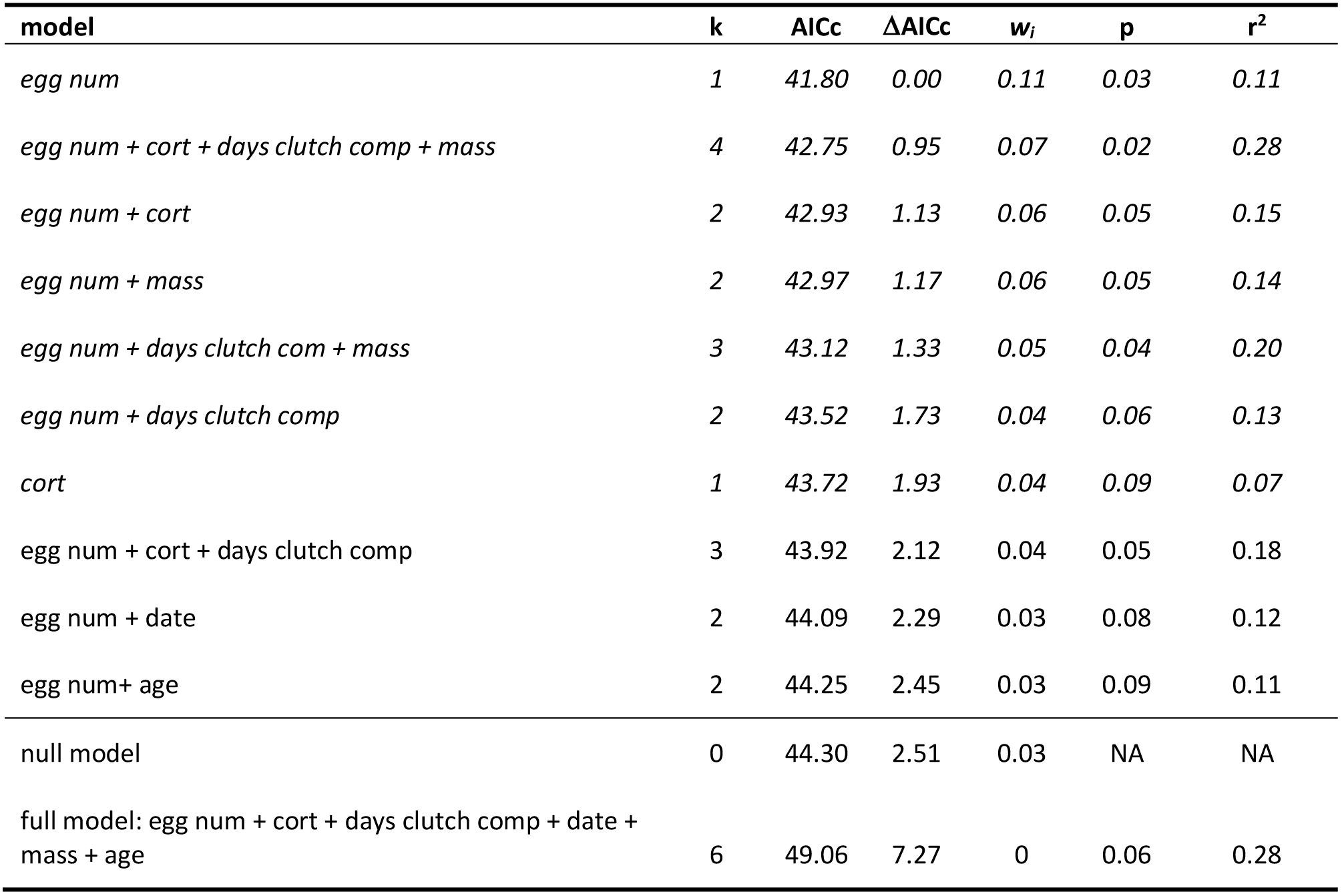
Generalized linear models predicting egg rejection at day 1 with AICc higher than the null model. k = number of fixed terms in the model, AICc = Akaike information criterion, ΔAICc = delta AICc between the focal model and the best model, w_i_ = Akaike weights, p = probability that the model explains more variation than the null model, r^2^ = McFadden’s pseudo r^2^ of model fit. Egg num: number of eggs at the time of parasitism; cort: baseline corticosterone adjusted for bleeding time, days clutch comp: number of days elapsed between clutch completion and parasitism, mass: female mass in g; age: second year or after second year age groups. Top seven models with ΔAICc < 2.0 are in italic font.

| model | k | AICc | $\Delta\text{AICc}$ | $w_i$ | p | $r^2$ |
| --- | --- | --- | --- | --- | --- | --- |
| *egg num* | 1 | 41.80 | 0.00 | 0.11 | 0.03 | 0.11 |
| *egg num + cort + days clutch comp + mass* | 4 | 42.75 | 0.95 | 0.07 | 0.02 | 0.28 |
| *egg num + cort* | 2 | 42.93 | 1.13 | 0.06 | 0.05 | 0.15 |
| *egg num + mass* | 2 | 42.97 | 1.17 | 0.06 | 0.05 | 0.14 |
| *egg num + days clutch com + mass* | 3 | 43.12 | 1.33 | 0.05 | 0.04 | 0.20 |
| *egg num + days clutch comp* | 2 | 43.52 | 1.73 | 0.04 | 0.06 | 0.13 |
| *cort* | 1 | 43.72 | 1.93 | 0.04 | 0.09 | 0.07 |
| egg num + cort + days clutch comp | 3 | 43.92 | 2.12 | 0.04 | 0.05 | 0.18 |
| egg num + date | 2 | 44.09 | 2.29 | 0.03 | 0.08 | 0.12 |
| egg num+ age | 2 | 44.25 | 2.45 | 0.03 | 0.09 | 0.11 |
| null model | 0 | 44.30 | 2.51 | 0.03 | NA | NA |
| full model: egg num + cort + days clutch comp + date + mass + age | 6 | 49.06 | 7.27 | 0 | 0.06 | 0.28 |

**Table 2.** GLM model predicting the probability of egg rejection at day 1, including all of the covariates present in the 7 top models; s.e. = standard error. Significant variables and p-values are italicized.

| covariate | estimate | s.e. | z | p |
| --- | --- | --- | --- | --- |
| (Intercept) | 0.61 | 0.48 | 1.25 | 0.21 |
| *egg number* | *-1.18* | *0.56* | *-2.11* | *0.04* |
| mass | -1.12 | 0.64 | -1.76 | 0.08 |
| corticosterone | -0.86 | 0.53 | -1.62 | 0.11 |
| days since clutch completion | -1.22 | 0.64 | -1.92 | 0.06 |

**Figure 2.**
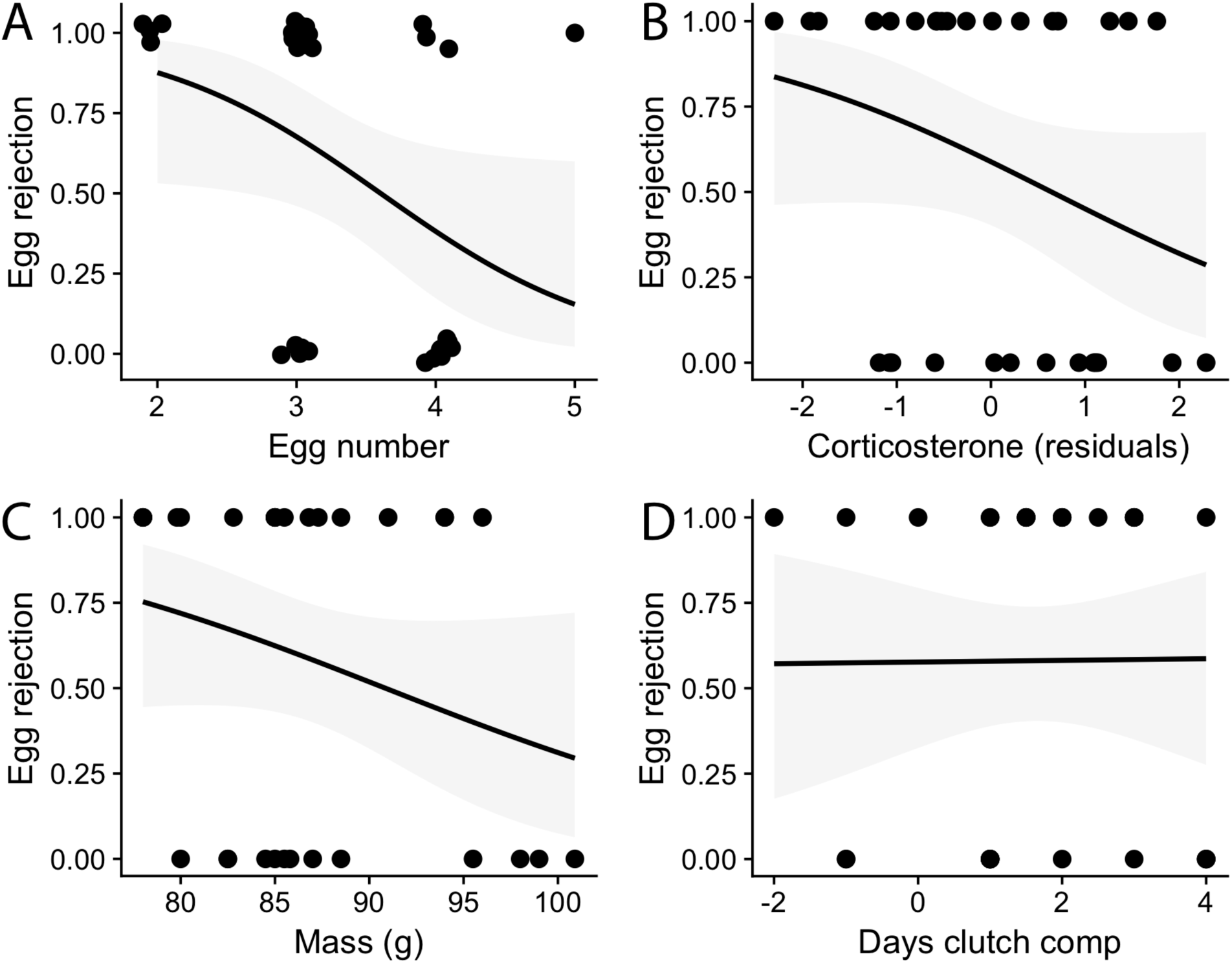
Relationship between the probability of egg rejection at day one (egg rejection) and (A) egg number at the time of parasitism, (B) baseline corticosterone levels (time-corrected residual levels), (C) mass, and (D) days between clutch completion and parasitism (“days clutch comp”). Grey shading indicates the 95% confidence intervals from the logistic regressions. Plots show relationship between raw data points. Data points at discrete x-values are jittered for visibility.

Despite the uncertainty of the composition of the best model, likelihood ratio tests showed that all except two of the top 7 models with ΔAICc < 2.0 were significantly better predictors of egg rejection behavior compared to the null model (Table 1). The number of eggs term was present in 6 out of 7 top models, with the only exception being a model that only contained corticosterone levels. Other predictors among the 7 models with ΔAICc < 2.0 - corticosterone, mass, and days since clutch completion - were present in 3 out of the 7 top models (Figure 2). The full model which included all of the covariates had a much lower AICc than the null model (Table 1).

Model averaging of all models with cumulative Akaike weight of 0.95 echoed the patterns of individual top models and suggested that the number of eggs at the time of the experimental parasitism was consistently the strongest predictor of egg rejection – birds with less eggs in the nest had a higher probability of egg rejection. This metric was present in 26 out of 45 models with a relative importance of 0.72 (Table 3). The other 3 predictors occurring in some of the top 7 models, had a weaker relationship to egg rejection: birds with lower mass (importance: 0.46, present in 24 out of 45 models), lower corticosterone levels (importance: 0.44, present in 21 out of 45 models), and birds experimentally parasitized earlier with respect to the clutch completion date (importance: 0.37, present in 18 of 45 models) were more likely to reject the model eggs.

**Table 3.** Averaged parameter estimates and the importance of covariates in explaining the probability of egg rejection on day 1; s.e. = adjusted standard errors, importance = sum of Akaike weights in all models that include the variable; CI = confidence interval.

| covariate | estimate (95% CIs) | s.e. | z | importance |
| --- | --- | --- | --- | --- |
| (Intercept) | 0.42 (-0.47, 1.31) | 0.45 | 0.93 | NA |
| egg number | -0.96 (-2.04, 0.11) | 0.55 | 1.76 | 0.72 |
| mass | -0.79 (-2.04, 0.46) | 0.64 | 1.24 | 0.46 |
| corticosterone | -0.60 (-1.53, 0.32) | 0.47 | 1.27 | 0.44 |
| days since clutch completion | -0.73 (-2.01, 0.55) | 0.65 | 1.12 | 0.37 |
| date | -0.09 (-1.36, 1.18) | 0.65 | 0.14 | 0.21 |
| age (older) | -0.17 (-2.06, 1.73) | 0.97 | 0.17 | 0.18 |

## Discussion

Rejection of parasitic eggs by hosts is one of the most widespread and effective defenses against avian brood parasites (Davies, 2000; Soler, 2017). However, different host species and, at an intraspecific level, different individuals show substantial variation in the propensity to reject foreign eggs from their nests (Alvarez, 1996; Brooke, Davies, & Noble, 1998; Bård G. Stokke, Moksnes, & RØskaft, 2005). Here we asked if cognitive, physiological, and life-history factors predict variation in egg rejection by a brood parasite host. Our correlational data set and model averaging approach showed that, in decreasing order of importance, among the most likely factors that may explain variation in model egg rejection by American Robins were the number of eggs in the nest at the time of the experimental parasitism, adult mass, the baseline corticosterone level, and the incubation stage relative to clutch completion: females with smaller clutches, lower mass, lower corticosterone, and those parasitized sooner after clutch completion were more likely to reject the model eggs. These relationships suggest that egg rejection is a complex behavior that is regulated by a diversity of factors.

The strongest and most consistent predictor of egg rejection in our models was clutch size, indicating that females with smaller clutches at the time of experimental parasitism were more likely to reject the model egg within a day compared to females with larger clutches. Because all of the females that were included in the analysis had completed laying their clutches at the time of experimental parasitism, this suggests that this relationship is not driven by variation in whether or not the clutch was completed, but by the final clutch size.

The effect of clutch size (and, thus, the parasite-to-host egg ratio) on egg rejection appears to vary between studies and species. On the one hand, multiple parasitism (higher ratio) has been shown to reduce the rejection of parasitic eggs (Bán et al., 2013; Manna et al., 2019; Moskát et al., 2009; M. Stevens et al., 2013) supporting the hypothesis that the ability of hosts to reject eggs is compromised because multiple eggs can be perceived as the most different when using a discordancy-based egg rejection mechanism (but see Lang, Bollinger, & Peer, 2014). On the other hand, irrespective of the extent of parasitism, greater absolute variation in egg coloration, associated with larger host clutches, has also been suggested to decrease the host’s ability to detect the foreign egg (Øein, Moksnes, & RØskaft, 1995). In support of this hypothesis, higher within-clutch variation due to more variable host eggs, or a bigger difference between the host and parasite egg colors, is associated with lower probability of egg rejection in mimetic parasite-host systems (Moskát et al., 2008; Stokke, Moksnes, Roskaft, Rudolfsen, & Honza, 1999), although it causes no change in egg rejection in non-mimetic parasite-host systems (Abernathy & Peer, 2014; Rebecca Croston & Hauber, 2015; Lang et al., 2014).

Yet another hypothesis that has not so far been explicitly addressed in host-brood parasite systems is that the proportion of parasite-to-host eggs in the nest affects the ability of hosts to perceive the parasitic egg due to the way animals respond to proportional differences in stimuli (Weber’s law; Akre, Farris, Lea, Page, & Ryan, 2011; Akre & Johnsen, 2014; Weber, 1978). Akre & Johnsen (2014) specifically proposed that in avian host-brood parasite systems, hosts may be better able to detect a parasitic egg if it is laid in a smaller clutch as opposed to a larger clutch. This is because in a smaller clutch, the addition of a parasitic egg causes a larger proportional increase compared to a large clutch. This, in turn, makes foreign eggs in smaller clutches easier to perceive and reject. The relationship between model egg rejection and clutch size in our study is consistent with the predictions of Weber’s law of proportional processing as applied to avian host-parasite systems (Akre & Johnsen, 2014). Importantly, however, because robins do not reject mimetic eggs (Luro et al., 2018; Rothstein, 1982), the relationship between clutch size and egg rejection in robins cannot be explained by a proportional increase in the clutch size alone (irrespective of color). Instead, a larger relative increase in the clutch size may affect the color threshold at which eggs are rejected (Abolins-Abols et al., 2019; Hanley et al., 2017, 2019). We suggest that proportional processing may therefore constitute an important aspect of the cognitive and perceptual processes, which, in concert, regulate egg rejection.

Female mass was negatively related to the rejection of model eggs in our study, indicating that heavier females either had higher thresholds for rejection of model eggs or showed, on average, higher maternal motivation to respond affiliatively towards (all) eggs. Mass has been associated with egg rejection in inter-specific comparisons (Peer & Sealy, 2004), where larger species are more likely to reject parasitic eggs. At an intraspecific level, heavier females may have higher expected future reproductive success (Abolins-Abols & Ketterson, 2017; Blums, Clark, & Mednis, 2002), possibly leading to a reduced sensitivity to parasitic stimuli. More focal experimental work will be needed to assess the mechanisms behind this association.

We also found evidence that females with lower corticosterone levels rejected the model eggs more rapidly, although the support for this relationship was statistically weak. This finding contradicts our prediction that, because higher corticosterone levels may lead to suppression of maternal attachment to eggs, robin females with higher baseline corticosterone levels should be more likely to reject the model eggs (Abolins-Abols & Hauber, 2018). However, we consider the alternative, namely that corticosterone may increase the motivation to care for eggs, unlikely. Instead, we argue that corticosterone levels may covary with or influence other phenotypic, sensory, and/or cognitive traits that are linked to egg perception, attention and, indirectly, to egg rejection propensity. Experimental tests with manipulated circulating hormone levels will be needed to assess these alternatives fully.

Finally, we found support for the pattern that females that experienced experimental parasitism earlier into the incubation stage were more likely to reject the model eggs. This agrees with similar findings in other avian host-parasite systems, where females are more likely to reject eggs earlier into the incubation stage of nesting (Moskát, 2005). This finding is consistent with at least two alternative hypotheses. First, because brood parasites target hosts nests during laying to ensure that the parasitic chicks hatch earlier or at the same time as host chicks, hosts should be selected to respond more selectively towards brood parasite eggs earlier in laying/incubation (Moskát, 2005). Alternatively, hosts may be less willing to eject any eggs later in incubation as a consequence of increasing general maternal motivation to care for advanced eggs or nestlings (Knight & Temple, 1986; Montgomerie & Weatherhead, 1988). Proximally, both of these alternative motivations – decrease in agonistic responses against brood parasitic stimuli, and an increase in general maternal motivation – may be mediated by hormones that regulate maternal care and motivation in other organisms, such as prolactin, estradiol, and progesterone (Abolins-Abols & Hauber, 2018; Bridges, Numan, Ronsheim, Mann, & Lupini, 1990; Cheng & Silver, 1975). Future studies should therefore also address these potential endocrine mediators of egg rejection observationally and experimentally.

In summary, we show that predicting egg rejection by American Robins requires complex models, and that this behavior is likely regulated by the perceptual environment, life history, as well as physiological factors. These factors may contribute to both plastic as well as individually consistent variation in egg rejection, although further studies are needed to experimentally establish the precise causal links between these variables and hot behavior. Despite our extensive efforts to include a diversity of proximate factors to understand variation in egg rejection, our best models were able to explain only 28% of variation in the rejection behavior. Research to assess what other factors underlie and cause variation in this ecologically and evolutionarily indispensable behavior therefore remains a potentially fruitful field.

## Ethical approval

All applicable national and institutional guidelines for the care and use of animals were followed. All procedures were approved by the University of Illinois IACUC (protocol #17259) and state and federal agencies.

