## Supplementary material for "Proximate Predictors of Variation in Egg Rejection Behavior by Hosts of Avian Brood Parasites": Figure S1

**Figure S1**. The relationship between the blood collection start time (in seconds from the moment of capture) and corticosterone levels (ng/mL). Blue line indicates linear model fit, and the shaded grey area indicates 95% confidence interval for the line.

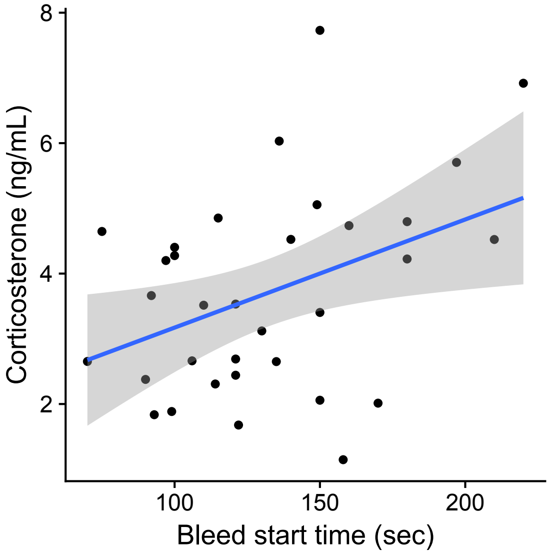
